# Structural basis for proton inhibition of the two-pore domain K^+^ channel TASK-1

**DOI:** 10.1101/2025.04.23.650279

**Authors:** Trevor Docter, Michelle S. Reid, Nicolas S. Lee, Stephen G. Brohawn

## Abstract

TASK-1 is a pH-sensitive two-pore domain K^+^ channel inhibited by external protons. TASK-1 is widely expressed, including in cardiomyocytes and chemosensitive neurons in the brain. Altered channel expression and activity are implicated in diseases including atrial fibrillation, pulmonary arterial hypertension, and developmental delay with sleep apnea. Structures of TASK-1 have been captured with an open external and closed internal gate. The structural basis for external proton inhibition and potential for communication between inner and outer gates are incompletely understood. Here, we determine a cryo-EM structure of TASK-1 at low pH with a closed extracellular gate and assess pH sensitivity of mutated channels. Proton inhibition of TASK-1 involves a C-type selectivity filter gate similar to TASK-3, but distinct from other proton-inhibited K2Ps TASK-2 and TWIK-1. Protonation of an extracellular histidine leads to formation of a hydrophobic seal above the selectivity filter and dilation of the outer two K^+^ coordination sites to close the channel pore. We find that a pulmonary arterial hypertension-associated loss-of-function mutation near the outer gate increases proton sensitivity, while gain-of-function mutations near the inner X-gate and intracellular cavity reduce proton sensitivity. These data reveal the structural basis for extracellular pH-gating of TASK-1, suggest allosteric communication between the inner and outer gates, and illustrate differences in C-type gating among related K^+^ channels.

## Introduction

Two-pore domain potassium (K2P) channels are named for their unique architecture in which channels assemble as homo- or heterodimers with a single conduction path. Each subunit contains two pore domains, consisting of two pore helices and two selectivity filter segments. K2Ps contribute to the resting potential of cells through basal leak activity that drives membrane voltage towards the Nernst potential for K^+^ (E_K+_ ≈ -90 mV). In addition, K2P activity is modulated by diverse chemical and mechanical stimuli to regulate cellular electrical activity^1,2^.

TASK-1 (TWIK-related acid-sensitive potassium channel 1, encoded by *KCNK3*) is a pH-gated K2P inhibited by extracellular protons (midpoint activation pH_1/2_ ∼ 7.3)^3,4^. TASK-1 is broadly expressed including in cardiomyocytes, vascular smooth muscle cells, and chemosensitive and other neurons in the brain^5–12^. TASK-1 is involved in the regulation of heart rate^6,7^, pulmonary arterial tone^13–16^, breathing control^17^, and response to volatile anesthetics^12^.

Loss-of-function mutations in TASK-1 result in inherited pulmonary arterial hypertension^16^, while gain-of-function mutations cause developmental delay with sleep apnea (DDSA)^17^. Channel upregulation is associated with atrial fibrillation^18^.

Structures of TASK-1 in detergent micelles and high [K^+^] (200 mM) have been determined at pH 8.5 (by X-ray crystallography) and pH 7.5 (by cryo-EM)^12,19^. They show that, like other K2Ps^20–23^, TASK-1 is a domain-swapped homodimer with four transmembrane helices per monomer, an extracellular helical cap, a selectivity filter, and an inner channel cavity. Both structures show an open extracellular gate and conductive selectivity filter conformation. The most striking difference between the two structures is a shift in the proton-sensing residue H98 from behind the selectivity filter at pH 8.5 up towards the extracellular solution at pH 7.5. Both structures also show a closed cytosolic X-gate formed by a ^243^VLRFMT^248^ motif near the C-terminal end of the fourth transmembrane domain (TM), raising questions about how this gate opens and whether it communicates with the extracellular pH gate.

TASK-3 is a related pH-gated K2P channel inhibited by protons with a pH_1/2_ ∼6.7^24^. Cryo-EM structures of TASK-3 in lipid nanodiscs were recently reported at pH 7.5 and 6.0 in high [K^+^] (200 mM), as well as pH 7.5 and 6.0 in low [K^+^] (5 mM)^25^, providing insight into the mechanism of channel gating by extracellular protons. At low pH and low K^+^, TASK-3 is closed on the extracellular side and adopts a nonconductive (“C-type inhibited”) selectivity filter conformation. Protonation of the pH sensing H98 residue results in a dilation of the selectivity filter at K^+^ coordination sites S1 and S2, accompanied by movement of selectivity filter residues Y96 and F202 to create a hydrophobic constriction above S1. Deprotonation of H98 removes the hydrophobic block and stabilizes a conductive filter conformation. Like TASK-1, TASK-3 adopts a closed cytosolic X-gate in all structures determined. Notably, pH regulation of two other K2Ps (TASK-2 and TWIK-1) involves completely distinct extracellular proton sensing and selectivity filter gating mechanisms^21,23^.

Here, we address two questions that arise from these results. First, what is the structural basis for proton inhibition of TASK-1? Second, is there communication between intracellular X-gate and extracellular selectivity filter gates? We determine a cryo-EM structure of TASK-1 in lipid nanodiscs at low pH and find proton inhibition involves a C-type selectivity filter gate similar to TASK-3, but distinct from those in TASK-2 and TWIK-1. Functional analysis demonstrates the intracellular cavity and X-gate tune the response of TASK-1 to extracellular protons and that disease-causing mutations impact pH sensitivity.

## Results

We reconstituted mouse TASK-1 in MSP1E3D1 lipid nanodiscs at pH 6.0 and 100 mM K^+^, conditions under which the channel is expected to be closed by extracellular protons^26^ (Fig. 1). We determined its structure by cryo-EM to an overall resolution of 3.3 Å, with higher local resolution for the core of the channel including around the selectivity filter (Figs. 1A-B, S1, S2, and Table 1). In contrast to previous TASK-1 structures^12,19^, the channel is closed on the extracellular side with a constriction above the filter that is too small for K^+^ permeation (Fig. 1C). The channel is also closed on the intracellular side below the channel vestibule by a constricted intracellular X-gate.

**Table 1.**
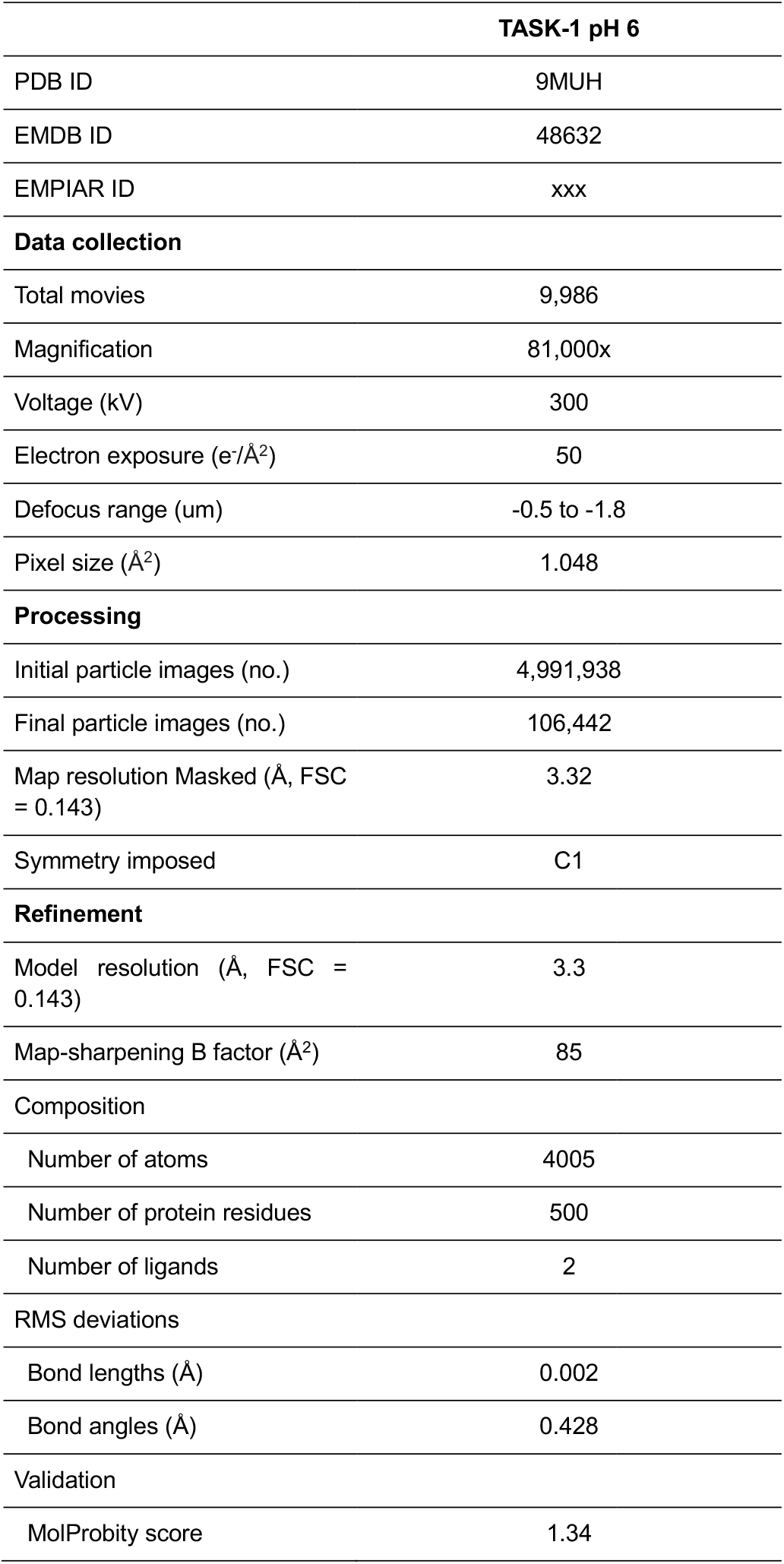
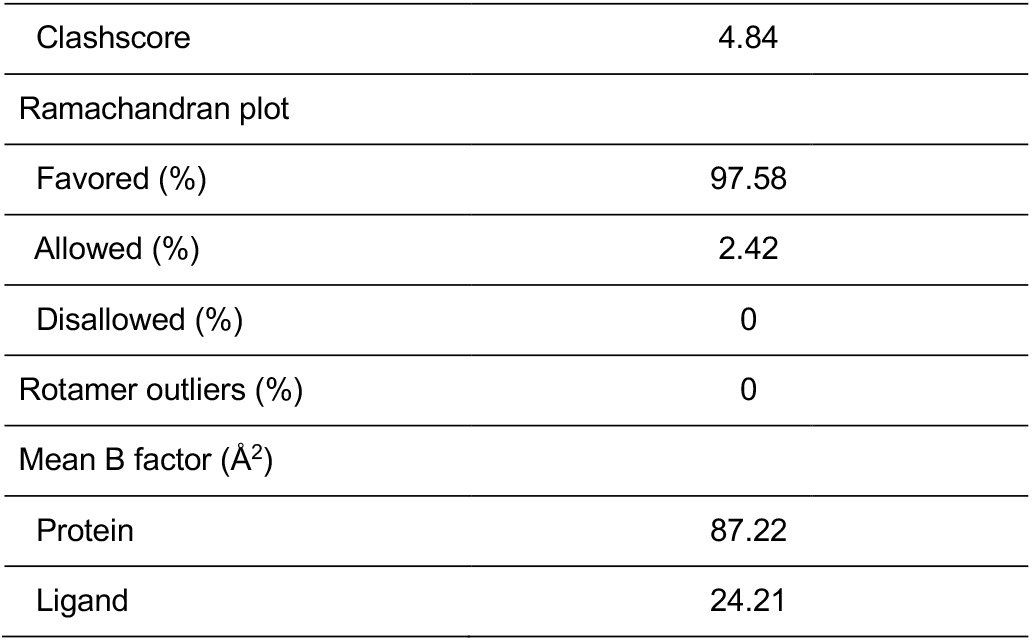
Cryo-EM data collection, refinement, and validation statistics.

**Figure 1.**
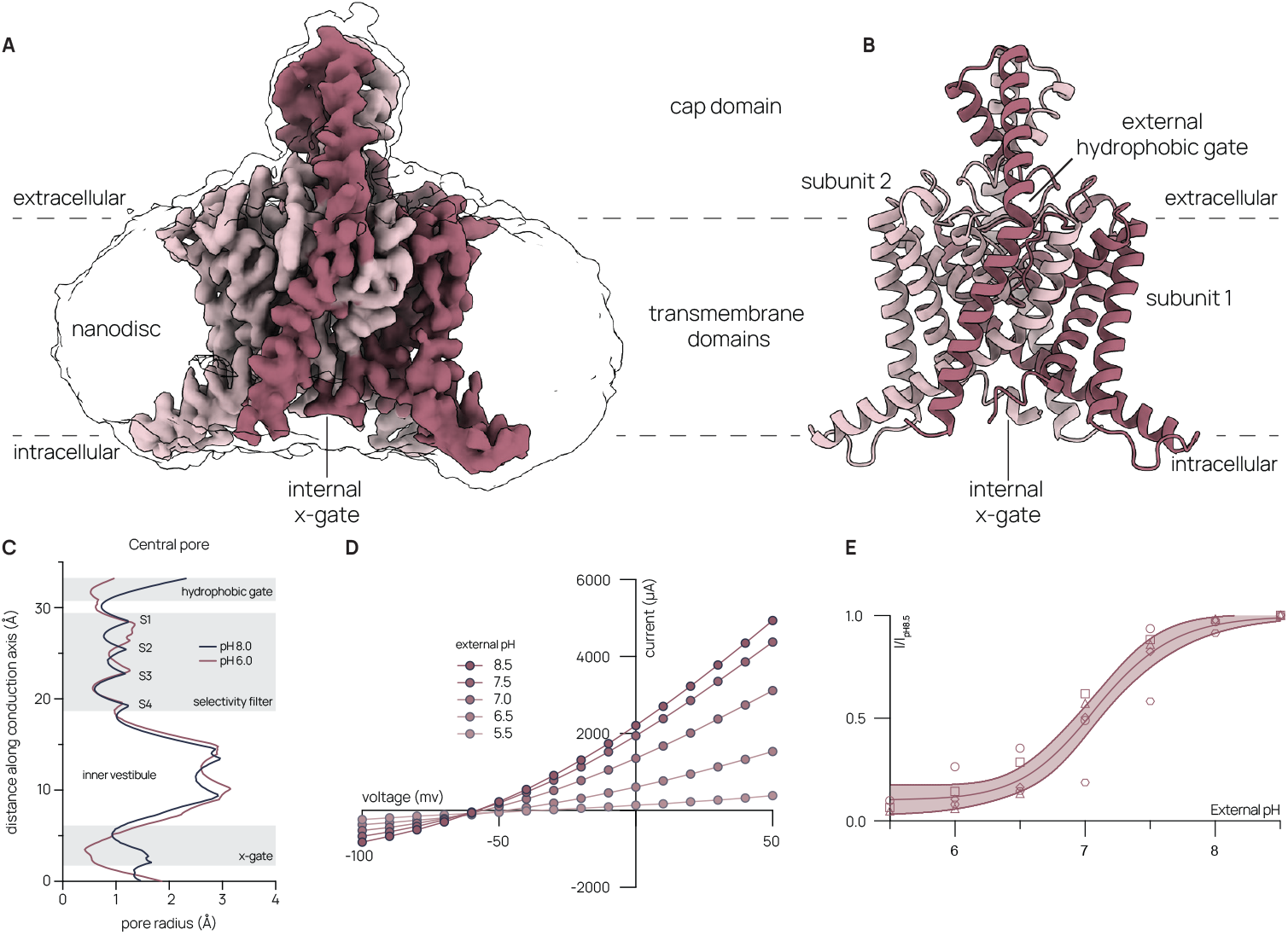
Cryo-EM structure of TASK-1 at pH 6.0 at 100 mM KCl. (A) 3.3 Å resolution cryo-EM map of Mus musculus TASK-1 at pH 6.0 in 100 mM KCl. TASK-1 subunits are shown in maroon and rose. Unsharpened map is shown transparent at low contour. (B) Overall model as shown from the membrane plane. (C) Comparison of pore radii between TASK-1 at pH 6.0 and 8.5 (PDB: 9MUH, this paper, and 6RV2 respectively^12^). (D) Representative current-voltage relationships from TASK-1 expressing X. laevis oocytes at variable external pH values. (E) TASK-1 dose-response curve to external pH measured at 0 mV. Data from individual cells are plotted with different shapes. Boltzmann fit with 95% confidence interval is shown (pH_1/2_ = 7.09 ± 0.03, R2 = 0.94, mean ± sem, n = 5 eggs).

Comparison of TASK-1 structures reveals how conformational changes around the extracellular side of the selectivity filter (residues G95-P101 and G201-A208) gate the channel closed at low pH (Figs. 1C, 2). In previously reported conductive filter conformations^12,19^, backbone carbonyl oxygens from Y96 and F202 coordinate K^+^ ions in site 1 (S1) while their side chains project away from the conduction axis to pack against pore and transmembrane helices. At low pH, the side chains of Y96 and F202 rotate ∼180º degrees upward, parallel to the K^+^ conduction axis, generating a hydrophobic constriction above the mouth of the filter that blocks K^+^ access to the pore (Fig. 2B-E). The transition is reminiscent of how one folds a cardboard box closed in the absence of tape. The movement involves dramatic rearrangement of S1 and S2 K^+^ coordinating carbonyls from Y96, F202, and G201 (by ∼180º. 180º, and ∼90º, respectively), dilating and altering the chemical environment at these sites. Consistently, density for K^+^ ions is absent in sites S1 and S2, but is present in sites S3 and S4, which adopt the same structure at high and low pH (Fig. 2A). We conclude TASK-1 adopts a non-conductive conformation at low pH through a C-type selectivity filter gate that involves hydrophobic block and disrupted K^+^ coordination.

**Figure 2.**
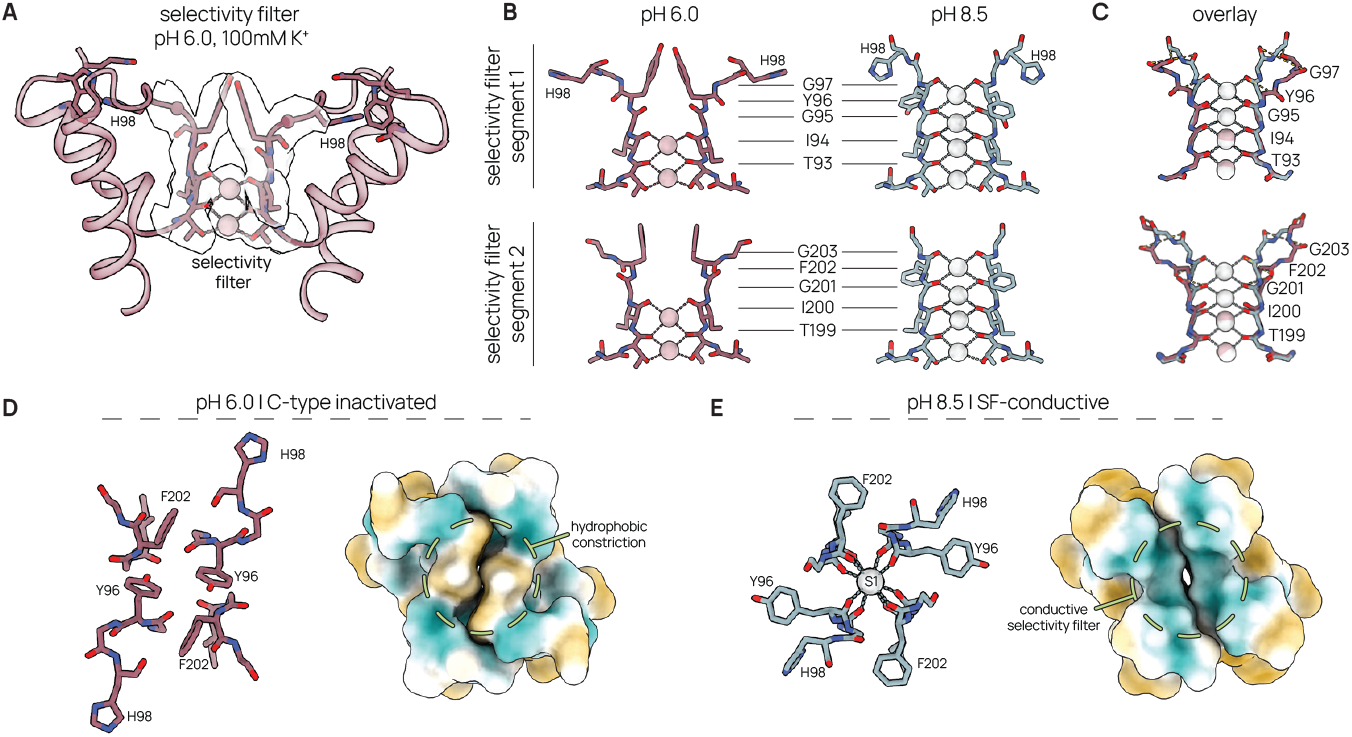
Structural basis for C-type inhibition of TASK-1 at the selectivity filter. (A) Model of pore domain 1 with transparent cryo-EM map density carved around selectivity filter and K^+^ coordination sites. (B) Comparison of the TASK-1 selectivity filter at pH 6.0 and pH 8.5 (C) overlay of SF1 and SF2 at pH 6.0 and 8.5 (PDB: 9MUH, this paper, and 6RV2, respectively^12^). (D), (E) top view of selectivity filter model (left) and hydrophobic surface (right) for C-type inhibited (D) and conductive (E) channels.

How are external protons sensed to trigger C-type gating? H98 is the principal proton sensor in TASK-1 channels^25,26^. At pH 8.5, when H98 is expected to be deprotonated, the imidazole side chain projects along the extracellular surface of the channel behind the selectivity filter (Fig. 2D-E). Hydrogen bonds between H98, two water molecules, and residues Q77, G89 and T89 stabilize the selectivity filter in a conductive state (Fig. 3A-C). At pH 7.5, H98 is present in a mixture of this conformation and one in which the side chain rotates out towards the extracellular solution^19^. At pH 6.0, when H98 is expected to be protonated, it rotates ∼90º and moves towards pore helix 1. Here, H98 packs between Q77 and G82, displacing a water molecule seen at pH 8.5, to form a cation–π interaction with W78 (imidazole nitrogen -tryptophan benzene distance ∼4.3 Å) (Fig. 3A). Movement of H98 requires rearrangement of Y96–A99 at the top of the first selectivity filter segment (Fig. 3A-C, 2B-E). G97 is pulled laterally away from the channel conduction axis to fill the space evacuated by H98 at pH 8.5. Y96 in turn flips up to accommodate the backbone reorganization. Concomitant movement of G201-G203 at the top of the second selectivity filter segment is required to a prevent a steric clash with Y96. G201 flips outward and F202 flips upward, resulting in a hydrophobic constriction above, and dilation of upper sites within, the selectivity filter.

**Figure 3.**
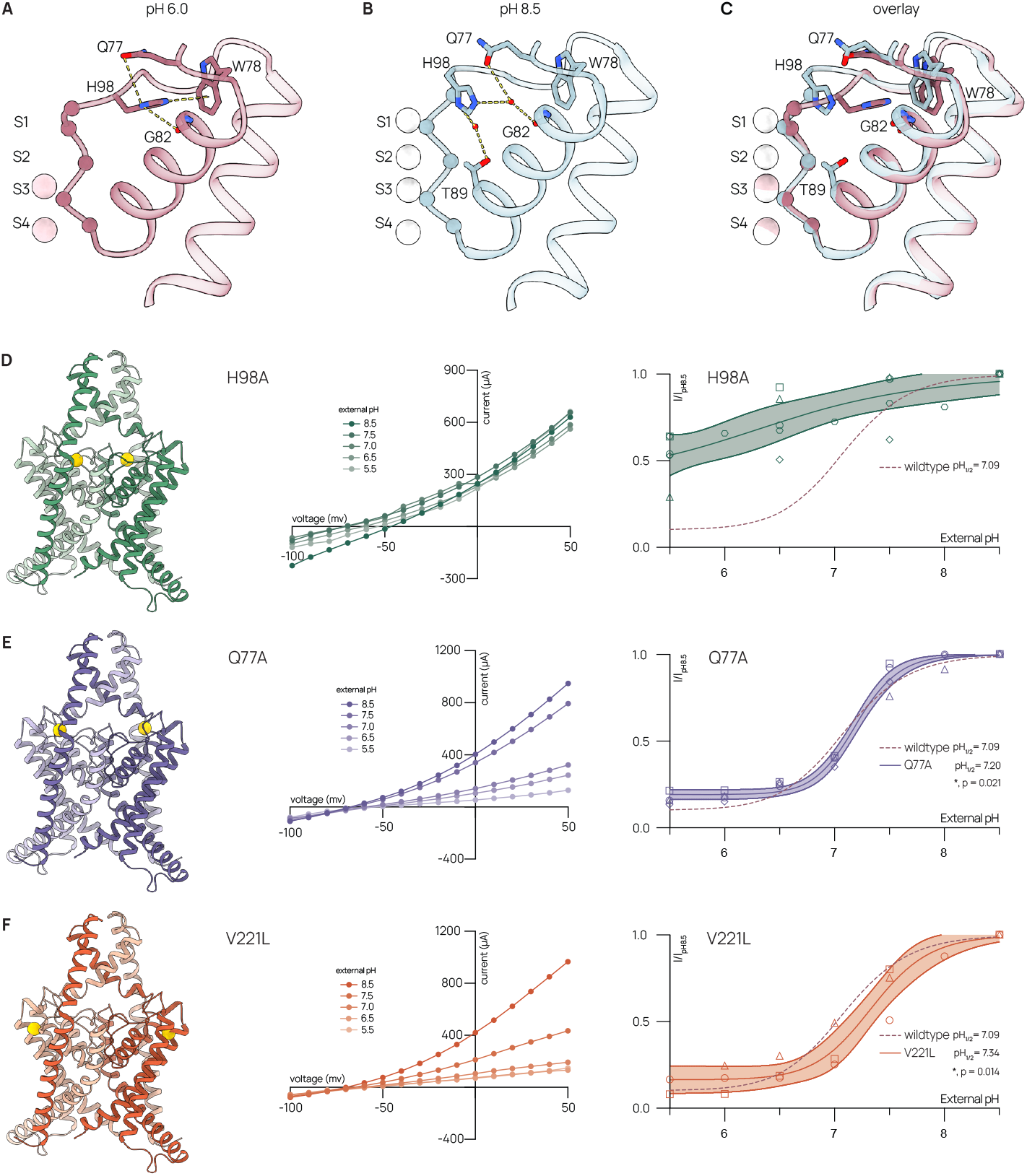
Structural basis for proton sensing in TASK-1. (A) Interaction network of H98 at pH 6.0 (rose) and (B) at pH 8.5 (navy). Waters are shown in red (PDB = 6RV2). (C) Overlay of structures at pH 6.0 (rose) and pH 8.5 (navy). Inhibition of (D) TASK-1_H98A_, (E) TASK-1_Q77A_, and (F) TRAAK_V221L_ by external protons (H98A: pH_1/2_ = 6.45 ± 0.35, R^2^ = 0.65, Q77A: pH_1/2_ = 7.20 ± 0.01, R^2^ = 0.99, V221L: pH_1/2_ = 7.34 ± 0.04, R^2^ = 0.94, (mean ± sem, n = 5, 4, 3 eggs, respectively). Structural models viewed from the membrane plane with mutated residue highlighted in yellow, representative current-voltage relationship, and dose-response curve at 0 mV shown from left to right. Data from individual cells are plotted with different shapes. Boltzmann fit to wild-type TASK-1 data shown with dotted maroon line.

To test this structural model for gating, we recorded activity of TASK-1 mutants expressed in *X. Laevis* oocytes using two-electrode voltage clamp while varying external pH. Similar to previous reports, we find wildtype TASK-1 is 90% inhibited at external pH 5.5 and reaches near maximal activation at external pH 8.5 with a pH_1/2_ = 7.09 ± 0.03 (Fig. 1D, E, Table 2)^25^. Mutation of the principal proton sensor H98A resulted in marked loss of pH sensitivity (Fig. 3D). A W78A mutation did not produce currents, suggesting a role for this residue in channel folding, trafficking, and/or opening. Q77A had modest, but significant, alterations in channel function, exhibiting more switch-like behavior around a right-shifted pH_50_ (7.20 ± 0.01) and higher residual activity at acidic pH values (Fig. 3E, Table 2). These results are consistent with interactions between Q77 and H98 stabilizing the closed state of TASK-1.

**Table 2.** Electrophysiological data summary.

| **Construct** | ***pH*<sub>1/2</sub>** | ***pH*<sub>1/2</sub>**<br>***p* value** | ***SEM*<sub>*pH*1/2</sub>** | ***Hill slope*** | ***SEM*<sub>*HS*</sub>** | ***Hill slope***<br>***p* value** | ***n*** | ***Fit R*<sup>2</sup>** |
| --- | --- | --- | --- | --- | --- | --- | --- | --- |
| Wildtype | 7.09 | - | 0.03 | -1.44 | 0.10 | - | 5 | 0.94 |
| H98A | 6.45 | - | 0.35 | -0.53 | 0.15 | - | 5 | 0.65 |
| Q77A | 7.20 | 0.021* | 0.01 | -2.21 | 0.10 | 0.003** | 4 | 0.99 |
| V221L | 7.34 | 0.014* | 0.04 | -1.62 | 0.21 | ns | 3 | 0.94 |
| L122N | 6.51 | 0.0023** | 0.06 | -1.24 | 0.17 | ns | 4 | 0.89 |
| L244A | 6.95 | 0.009** | 0.01 | -1.57 | 0.07 | ns | 6 | 0.97 |

Comparisons made using Brown-Forsythe and Welch ANOVA tests.

We next evaluated reported loss-of-function TASK-1 mutations implicated in pulmonary arterial hypertension ^16,19^. Mutations of glycines at the top of selectivity filter segments 1 or 2 (G97R, G203D and G203R) did not produce currents. However, a V221L mutant produced a significant right-shift of the Boltzmann fit toward more basic pH values (pH_1/2_ = 7.34 ± 0.04) (Fig. 3F, Table 2). This effect, which would result in reduced activity under physiological conditions, is consistent with our structural model. V221 sits near the top of TM4 where it forms part of the pocket for H98 in the protonated state and interacts with the linker between selectivity filter segment 2 and TM4 (Fig. S3). Steric clash between leucine and the linker region in the high pH state likely destabilizes the open state.

We observe a closed intracellular X-gate (Fig. 4A,B) like previously reported TASK-1 structures. Interaction between residues L244-T248 from each subunit seals the inner cavity under the selectivity filter from the cytoplasm (Figs. 1C, 4A,B). An L244A mutation that is predicted to disrupt X-gate formation has been shown to increase channel activity^12,17^, but its effect on pH sensitivity has not been reported. We found L244A results in a modest, but significant, leftward shift of the pH activation curve (pH_1/2_ = 6.95 ± 0.01) (Fig. 4C, Table 2). This may contribute to increased activity of L244A and raises the possibility of allosteric communication between channel gates.

**Figure 4.**
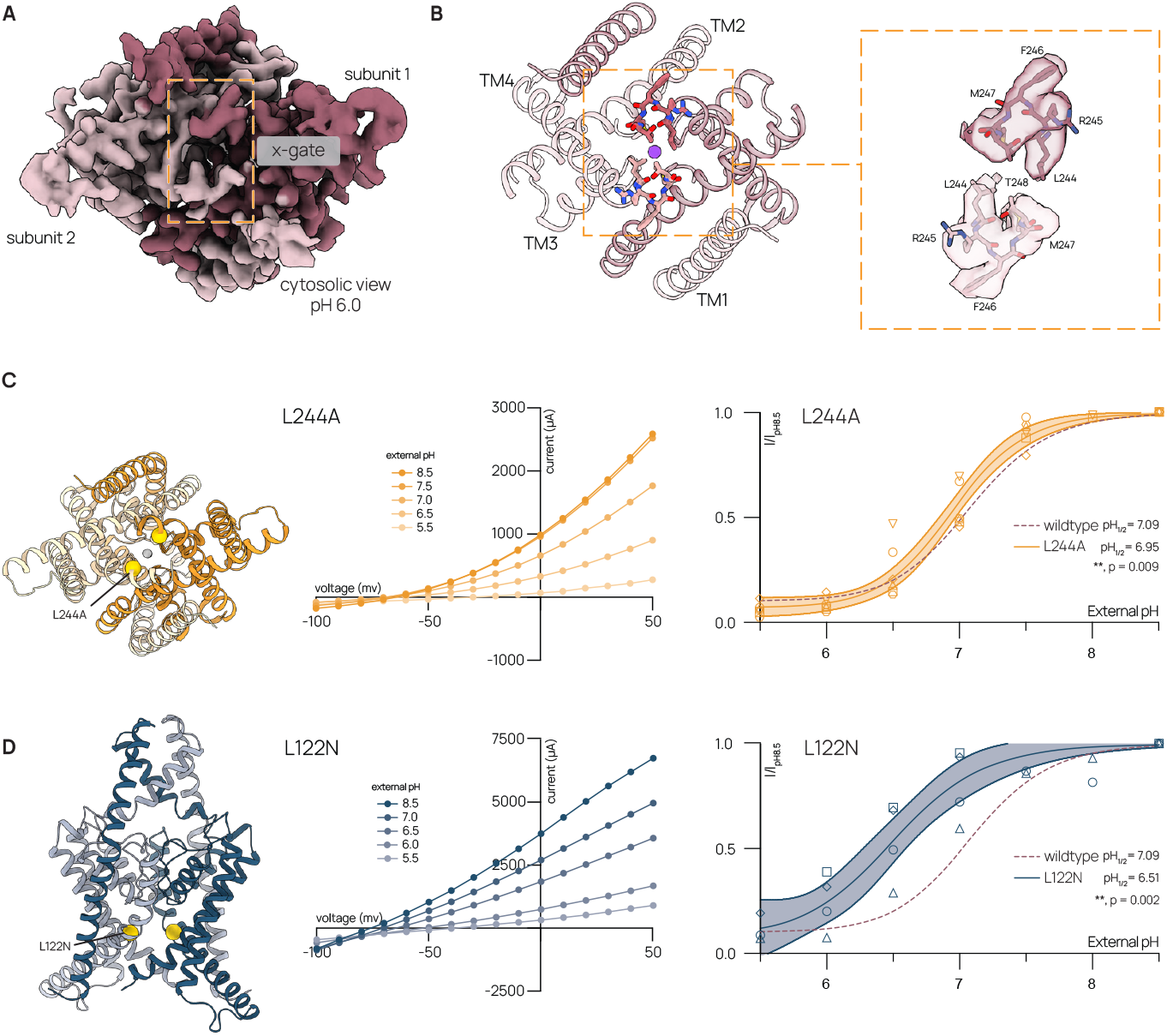
Intracellular gain of function mutations reduce sensitivity to external protons. (A) Cytosolic view of the cryo-EM map of TASK-1 with internal X-gate highlighted. (B) TASK-1 model at pH 6.0 (left) with zoom in on the residues forming the internal X-gate (LRFMT) and the corresponding carved cryo-EM density. Inhibition of (C) TASK-1_L244A_ and (D) TASK-1_L122N_ by external protons (L244A: pH_1/2_ = 6.95 ± 0.01, R^2^ = 0.97, L122N: pH_1/2_ = 6.51 ± 0.06, R^2^ = 0.89 (mean ± sem, n = 6 and 4 eggs, respectively). Structural models viewed from the (C) cytosolic side or (D) membrane plane with mutated residue highlighted in yellow, representative current-voltage relationship, and dose-response curve at 0 mV shown from left to right. Data from individual cells are plotted with different shapes. Boltzmann fit to wild-type TASK-1 data shown with dotted maroon line.

To further explore this idea, we tested pH sensitivity of channels with mutations in the inner cavity above the X-gate. L122 on TM2 faces the cavity and mutations of this residue increase channel activity. L122P and L122V are reported gain of function mutations in DDSA patients ^17^ and L122N was characterized in a study showing all studied K2Ps can be activated by mutations that increase polarity at this position^27^. Whether these mutations modulate pH sensitivity in TASK-1 is not known. Strikingly, we found L122N resulted in a large reduction in proton sensitivity (pH_1/2_ = 6.51 ± 0.06) (Fig. 4D, Table 2). We conclude that gain of function from intracellular mutations is at least partially explained by altered extracellular pH sensitivity and that the internal X-gate and cavity of TASK-1 interact allosterically with the external pH gate.

## Discussion

In this study, we present a structure of the pH-sensitive channel TASK-1 in a fully closed state that illustrates the mechanism for proton inhibition through a C-type selectivity filter gate. A model for pH gating is presented in Fig. 5. Protonation of H98 results in two major conformational changes that gate the channel closed: (1) formation of a hydrophobic seal by Y98 and F202 and (2) disruption of the size and chemistry of K+ coordination sites S1 and S2.

**Figure 5.**
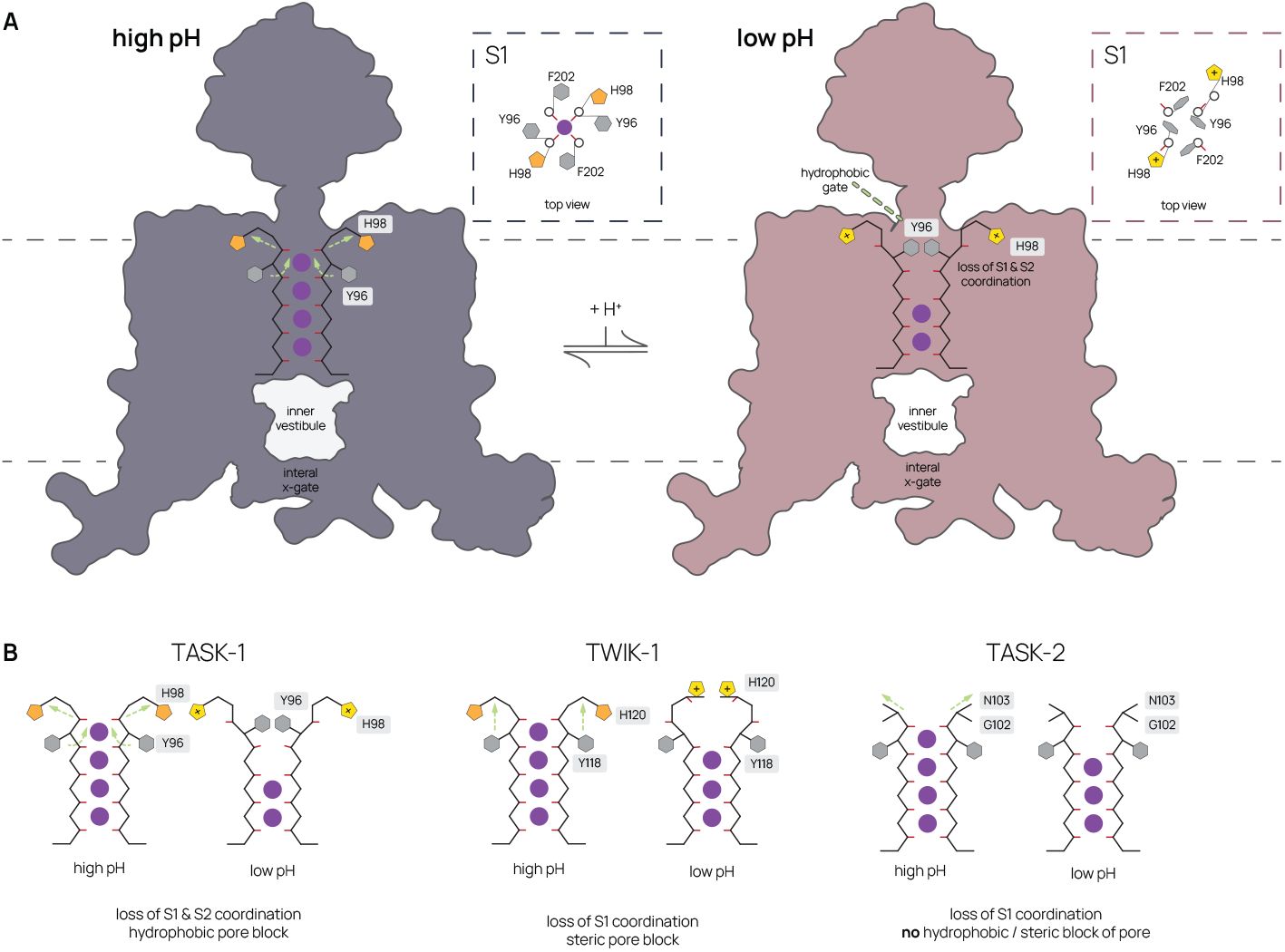
Structural model for TASK-1 gating in response to changes in external pH. (A) Model for selectivity filter gating of TASK-1 by external pH view from the membrane plane. TASK-1 adopts a conductive selectivity filter at high pH (purple, left). Protonation of H98 at low pH results in a conformational change in the selectivity filter that disrupts K+ coordination sites S1 and S2 and generates a hydrophobic seal above the selectivity filter (rose, right). The internal X-gate is predominantly closed under both conditions. Top view of the S1 site of the selectivity filter is shown in the boxed inset. Potassium ions are shown in purple. (B) Comparison of conformational changes to the selectivity filters of related pH-sensitive K2Ps TWIK-1 and TASK-2. Proton inhibition involves distinct C-type gating mechanisms.

Together, these rearrangements prevent ion access to the filter and conduction (Fig. 5A). At pH 8.5, deprotonated H98 is stabilized through a hydrogen bonding network between waters and residues Q77, G82, and T89. Upon protonation, H98 repositions into a pocket between Q77 and G82 to form a stabilizing by cation–π interaction with W78. Movement of H98 requires rearrangement of selectivity filter segments and subsequent linkers that form the hydrophobic gate and disrupted selectivity filter.

TASK-1 shares relatively high (54%) sequence homology with TASK-3. Like TASK-1, TASK-3 is widely expressed throughout the CNS and peripheral tissues^24,28–35^. TASK-1 and TASK-3 form homo-and hetero-dimeric channels physiologically^26,36^. Comparing structures of TASK-1 and TASK-3 show that pH gating is accomplished in a similar way^25^. Both channels are gated by conformational changes induced, at least in part, by extracellular protonation of H98 that results in hydrophobic pore block above the mouth of the channel and disruption of outer selectivity filter sites. The conserved nature of the conformational changes is relevant to the function of TASK-1/3 heteromers. Transition to the C-type inhibited state likely requires coordinated movement of all four selectivity filter regions. Protonation of H98 and associated conformational changes in one subunit is expected to create steric clashes with a second unprotonated subunit and thereby promote its protonation. This may explain positive cooperativity in homodimeric TASK-1 (Table 2) and the observation that heterodimers have a pH_1/2_ value close to the higher affinity TASK-1 subunit^4,24,26,36^.

Like TASK-1/-3, TWIK-1 and TASK-2 K2Ps are also inhibited by extracellular protons via selectivity filter gates, but this is accomplished through distinct mechanisms (Figs. 5B, S4A). The primary proton sensor in TWIK-1 is a histidine in the same position as TASK-1/-3 H98 and its protonation results in steric block and disrupted K^+^ coordination in the filter^23,25^. In TWIK-1 however, protonation results an electrostatic zipper-like seal above the filter formed by the histidine, threonine, and aspartic acid from opposing filters in contrast to the hydrophobic ring of phenylalanine and tyrosine side chains that seals TASK-1/-3 like a closed aperture (Fig. 5B)^23,25^. C-type gating of TASK-2 is accomplished by disruption of K^+^ binding in the filter without formation of a protein seal. The primary proton sensor in TASK-2 is an arginine at the top of TM4, more distally positioned from the filter than the histidines in TASK-1, TASK-3, and TWIK-1. Protonation of this arginine results in conformational changes that are relayed through the helical cap to disrupt geometry of K^+^ coordination sites S1 and S2^21^. This highlights how diverse structural mechanisms can underlie C-type gating of the selectivity filter, even in related channels that evolved respond to the same stimulus.

There remain open questions about the cytosolic gate in TASK-1 and TASK-3. All reported structures to date show a closed X-gate^12,19,25^(Figs. 1,4,5). How the X-gate is modulated is not known, but it is likely important for controlling TASK-1 activity and may explain low channel open probability^17^. Control by specific lipid binding to the channel has been postulated based on nearby cholesterol-like densities in structures^12,19,25^. We find that mutational disruption of the X-gate also impacts pH sensitivity, suggesting allosteric coupling between inner and outer gates.

Notably, mutation of a cavity-facing residue above the X-gate resulted in more pronounced reduced proton sensitivity. This suggests cavity mutation promotes states with conductive selectivity filters and further supports the notion that inner and outer gates operate synergistically to control channel activity. Whether cavity mutations in TASK-1 impact other aspects of channel gating remains to be determined. One possibility if that cavity mutations disrupt nearby X-gate formation directly. A second possibility is that cavity mutations disrupt potential lipid binding in the cavity and block of ion conduction. Lateral fenestrations connecting the cavity to the surrounding membrane are present in TASK-1(Fig. S4B), like other structurally characterized K2Ps^20–23,37,38^. Lipids are proposed to enter the fenestrations and block ion conduction in TWIK-1, TWIK-2, TRAAK, and TREK-1^20,22,23,37,38^. Mutations at the same position as TASK-1/3 L122 (on TM2 facing the cavity) activate channels across the K2P channel family^27^ and we showed the analogous mutation in TRAAK disrupts lipid block of the cavity to increase channel activity^20^. Evidence for lipids in TASK-1 or TASK-3 structures has not been reported, but ours and other structures may not be determined to resolutions sufficient for their clear identification. Further work is required to better understand the nature of inner gating and mechanisms for allosteric communication between inner and outer gates in TASK-1/-3 channels.

### Methods Electrophysiology

For two-electrode voltage clamp recording from *Xenopus* oocytes, the gene encoding full-length *Homo sapiens* TASK-1 (Uniprot O14649) was codon optimized for eukaryotic expression, synthesized (Genewiz), and cloned into a modified pGEMHE vector using XhoI and EcoR1 restriction sites. The transcribed messages encode *H. sapiens* TASK-1 amino acids 1– 394 with an additional three amino acids (SNS) at the C terminus. TASK-1 mutants were introduced by polymerase chain reaction (PCR). Linearized DNA was transcribed *in vitro* using T7 polymerase. Complementary RNA (0.1 to 50 ng for TASK-1 and mutants) in 50 nL H_2_O was injected into *Xenopus laevis* oocytes extracted from anesthetized frogs. Currents were recorded at 25 °C from whole oocytes 1 to 5 d after RNA injection. The pipette solution contained 3M KCl and the bath solution contained 150 mM KCl and 10 mM HEPES (pH = 7.1 with KOH).

Borosilicate glass pipettes were pulled to 2-5 MΩ resistance.

Currents were recorded at 25 °C using two-electrode voltage clamp (TEVC) from oocytes 2 days after mRNA injection. Pipette solution contained 3 M KCl. Bath solution contained ND96 (96 mM NaCl, 2 mM KCl, 1.8 mM CaCl_2_, 1 mM MgCl_2_ 2, 2.5 mM Na Pyruvate, 20 mM buffer) at a pH range of 5.5 to 8.5. MES was used to buffer to pH 5.5 – 6.5 and HEPES was used to buffer each solution to a pH of 7.0 – 8.5. Bath volume was kept at ∼300 uL during recording. Eggs were initially placed in the recording chamber in ND96 at pH 8.5 to confirm channel currents. Eggs were then perfused in the bath with solutions ranging from pH 8.0 to pH 5.5. Currents were recorded and low pass filtered at 2 kHz using an Axon Instruments Axoclamp 900A amplifier and digitized at 10 kHz with a Sutter Dendrite digitizer.

The effects of pH on channel activity were quantified by the reduction in channel current evoked at 0 mV during 200 ms long voltage families with steps ranging from -100 to +40 mV. Currents were normalized to the maximum current recorded at the highest pH experienced by the egg to facilitate comparisons across different cells. Dose response curves were four-parametric Boltzmann fits to aggregated data done at 0mV. For voltage-step recordings, current was measured at 0 mV from each recording. Statistical comparisons were conducted with ordinary one-way ANOVA, Brown-Forsythe and Welch ANOVA, and Tukey multiple-comparison tests using Graphpad Prism.

### TASK-1 expression and purification

A gene encoding *Mus musculus* TASK1 (Uniprot O35111) was codon-optimized for expression in *Pichia pastoris*, synthesized (Genewiz, Inc), and cloned into a modified pPICZ-B vector (Life Technologies Inc). The final construct is C-terminally truncated by 153 amino acids and expressed as a C-terminal PreScission protease-cleavable fusion to EGFP (mmTASK1 amino acids 1-256 – SNSLEVLFQGP – EGFP – 10xHis). Pme1 linearized plasmid was transformed into *P. pastoris* strain SMD1163 by electroporation and transformants were selected on YPDS plates with 1 mg/mL zeocin. For large-scale expression, overnight cultures of cells in YPD with 0.5 mg/mL Zeocin were added to BMGY and grown overnight at 30 °C to a final OD 600 ∼ 25. Cells were centrifuged at 8000 x g for10 min and added to BMMY at 27 °C to induce protein expression. Expression continued for ∼24-48 h. Cells were harvested and flash frozen in liquid nitrogen.

60g of cells expressing mmTASK-1 were disrupted by milling (Retsch model MM301) 5 times for 3 min at 25 Hz with chamber cooled in liquid nitrogen between cycles. All subsequent purification steps were carried out at 4°C. Milled cells were thawed in 100 mL of lysis buffer (50 mM TRIS, 150 mM KCl, 1mM EDTA pH 8, unless otherwise noted, the pH values for Tris buffers correspond to values at 4 °C). Protease inhibitors (Final Concentrations: E64 (1 μM), pepstatin A (1 μg/mL), soy trypsin inhibitor (10 μg/mL), benzamidine (1 mM), aprotinin (1 μg/mL), leupeptin (1μg/mL), AEBSF (1mM), PMSF (1mM)), benzonase (10µL) and DNAse (10 µL) were added to the lysis buffer immediately before use. The solution was sonicated and centrifuged at 150,000 x g for 45 minutes. The supernatant was discarded, and residual nucleic acid was removed from the top of the membrane pellet using DPBS. Membrane pellets were transferred to a Dounce homogenizer containing extraction buffer (50 mM TRIS, 150 mM KCl, 1 mM EDTA, 1.0 % n-Dodecyl-β-D-Maltopyranoside (DDM, Anatrace, Maumee, OH), 0.2% cholesteryl hemisuccinate (CHS, Anatrace, Maumee, OH) pH 8). A stock solution of 10% DDM, 2% CHS was dissolved and clarified by bath sonication in 200 mM Tris pH 8 prior to addition to buffer to the indicated final concentration. Membrane pellets were homogenized in 100 mL extraction buffer and this mixture then stirred at 4°C for 2 hours. The extraction was centrifuged at 33,000 x g for 45 minutes and the supernatant, containing solubilized membrane protein, was bound to 4 mL of Sepharose resin coupled to anti-GFP nanobody for 2 hours at 4°C. The resin was collected in a column and washed with 10 mL of buffer 1 (20 mM TRIS, 150 mM KCl, 1 mM EDTA, 0.025% DDM, 0.005% CHS, pH 8), 40 mL of buffer 2 (20 mM TRIS, 500 mM KCl, 1 mM EDTA, 0.025% DDM, 0.005% CHS, pH 8), and 10 mL of buffer 1. The resin was then resuspended in 6 mL of buffer 1 with 0.5 mg of PPX protease and rocked gently in the capped column overnight (∼ 12–14 h). Cleaved TASK-1 was eluted in buffer 1, spin concentrated to ∼1 mL with an Amicon Ultra spin concentrator (100 kDa cutoff, Millipore) and loaded onto a Superdex S200 increase column (GE Healthcare, Chicago, IL) on an NGC system (Bio-Rad, Hercules, CA) equilibrated in SEC buffer 1 (20 mM HEPES, 150 mM KCl, 1 mM EDTA, 0.025% DDM, pH 7.4). Peak fractions containing TASK-1 protein were collected and spin concentrated for nanodisc reconstitution.

TASK1 was reconstituted into MSPE3D1 nanodiscs with a 2:1:1 DOPE:POPS:POPC lipid mixture (molar ratio, Avanti, Alabaster, Alabama) at a final molar ratio of TASK1:MSP1D1:lipid of 1:5:250. Lipids in chloroform were mixed, dried under argon, washed with pentane, dried under argon, and dried under vacuum overnight in the dark. Dried lipids were rehydrated in buffer containing 20 mM Tris, 150 mM KCl, 1 mM EDTA, pH 8.0 and clarified by bath sonication. DDM was added to a final concentration of 8 mM. TASK-1 was mixed with lipids and incubated at 4 °C for 1 h before addition MSPE3D1 protein. After incubation for 30 minutes at 4 °C, 100 mg of Biobeads SM2 (Bio-Rad, USA) (prepared by sequential washing in methanol, water, and buffer and weighed damp following bulk liquid removal) was added and the mixture was rotated at 4 °C overnight. The sample was spun down to facilitate removal of solution from the Biobeads and the reconstituted channel was further purified on a Superdex 200 increase column run in SEC buffer 2 (20 mM MES, 100 mM KCl, 1 mM EDTA pH 6.0). The peak fractions were collected and spin concentrated (50 kDa MWCO) to 0.5 mg/mL for grid preparation.

### Cryo-electron microscopy

TASK-1 in nanodiscs at 0.5 mg/mL was centrifuged at 21,000 x g for 10 min at 4 °C. A 3 μL sample was applied to holey carbon 400 mesh R 1.2/1.3 gold grids with an additional 2 nm carbon coating (Quantifoil, Großlöbichau, Germany) that were freshly glow discharged for 30 seconds. Sample was incubated for 5 seconds at 4 °C and 100% humidity prior to blotting with Whatman #1 filter paper for 3 seconds at blot force 1 and plunge-freezing in liquid ethane cooled by liquid nitrogen using a FEI Mark IV Vitrobot (FEI/Thermo Scientific, USA).

Grids were clipped and stored in liquid nitrogen. Data was collected on a Titan Krios G3i electron microscope operated at 300 kV equipped with a Gatan BioQuantum Imaging Filter with a slit width of 20 eV. Dose-fractionated images (50 electrons per Å^2^ over 50 frames) were recorded on a K3 direct electron detector (Gatan) in super-resolution counting mode with pixel size of 0.524 Å. Eighteen movies were collected in a 3 × 3-hole pattern with two targets per hole around a central hole position using image shift. Defocus was varied from −0.5 to −1.8 μm using SerialEM^39^. See Table 1 for data collection statistics.

Motion correction was performed on 9,989 micrographs using UCSF MotionCor2 in RELION v4.1^40^. Contrast transfer function (CTF) parameters were fit with patch CTFFIND v4.1^41^. Laplacian-of-gaussian (LOG) auto-picking of particles was performed with a minimum particle size of 120 Å and a maximum particle size of 200 Å on 9,045 movies CTF fit to 7.0Å or better, yielding an initial set of 4,734,533 particles. These were extracted at a 288-pixel box size for two-dimensional (2D) classification. Iterative rounds of 2D and 3D classification were conducted in cryoSPARC and yielded a set of 60,956 particles that were used to train a Topaz model^42,43^. Topaz-trained particle-picking yielded 4,991,938 particles which were manually curated through iterative rounds of 2D classing. A 3-class ab-initio was performed and the particles that corresponded to the best map density were chosen. One additional round of 2D classification was performed. A non-uniform refinement (C1, 4 extra passes, 15Å initial resolution) yielded a 3.53 Å map from 152,997 particles (Fig. S3). Due to high anisotropy, side views were selected following 2D classification and used to train additional Topaz models to select for side views. This yielded a set of 24,796 particles for additional rounds of Topaz training.

24,796 particles were extracted from three separate sets of 3000 micrographs to ensure topaz training on all micrographs and particles that presented side views. We generated three separate Topaz models (Topaz model 1, model 2, and model 3). After training and extraction from all micrographs with CTF fits <7.0 Å, Topaz model 1 yielded 1,491,161 particles, model 2 yielded 1,979,628 particles, and model 3 yielded 1,522,112 particles. Each particle set was curated via iterative rounds of 2D classification specifically looking for improved side views. After final rounds of 2D classification, model 1 yielded 158,454 particles, model 2 yielded 164,803 particles, and model 3 yielded 169,354 particles. These three particle sets were combined with the 152,997 particles from the previous non-uniform refinement, and duplicate particles were removed yielding 411,693 particles.

These particles were further cleaned with two rounds of 2D classification and then used for a non-uniform refinement using the previously generated best map to confirm if additional side views had been found and there had been an improvement in observed anisotropy. We saw a dramatic improvement in view angles and map quality. We then performed a 2 class ab-initio (C1, 0 sim) and chose the map with the clearest protein signal leaving 219,074 particles. These particles were then moved back into RELION for Bayesian polishing.

Polished particles were subjected to two rounds of heterogenous refinement using the previous ab-initio map and three junk classes, yielding 102,442 particles. Ab-initio and non-uniform refinement of these particles (C1, 4 extra passes, 15 Å initial resolution) resulted in a map with 3.29 Å overall resolution. Final FSC validation of this map confirmed a final resolution of 3.32 Å.

The final non-uniformed local refined and sharpened map from cryoSPARC was used for modeling. An initial apo TASK-1 model (PDB: 6RV2^12^) was rigid body fit to the density in Phenix^44^. Model building was performed iteratively with manual adjustment in Coot^45^, global real space refinement in Phenix^46^, and geometry assessment in Molprobity^47^. Figures were generated with a map sharpened in DeepEMHancer^47^.

## Acknowledgements

We thank D. Toso, P. Tobias, and R. Thakkar at the Cal-Cryo Electron Microscopy Facility for help with data collection. This work was supported by the National Institute of General Medical Sciences grant GM145869.

## Author Contributions

M.S.R., T.A.D., and S.G.B. conceived of the project. T.A.D. and N.L. performed electrophysiological experiments and analyzed data. M.S.R. performed protein expression, protein purification, cryo-EM sample preparation, and cryo-EM data collection. T.A.D performed cryo-EM data processing. T.A.D. and S.G.B. performed structure modeling and refinement.

T.A.D. and S.G.B. wrote the manuscript. S.G.B. secured funding.

## Figure Legends

**Figure S1.**
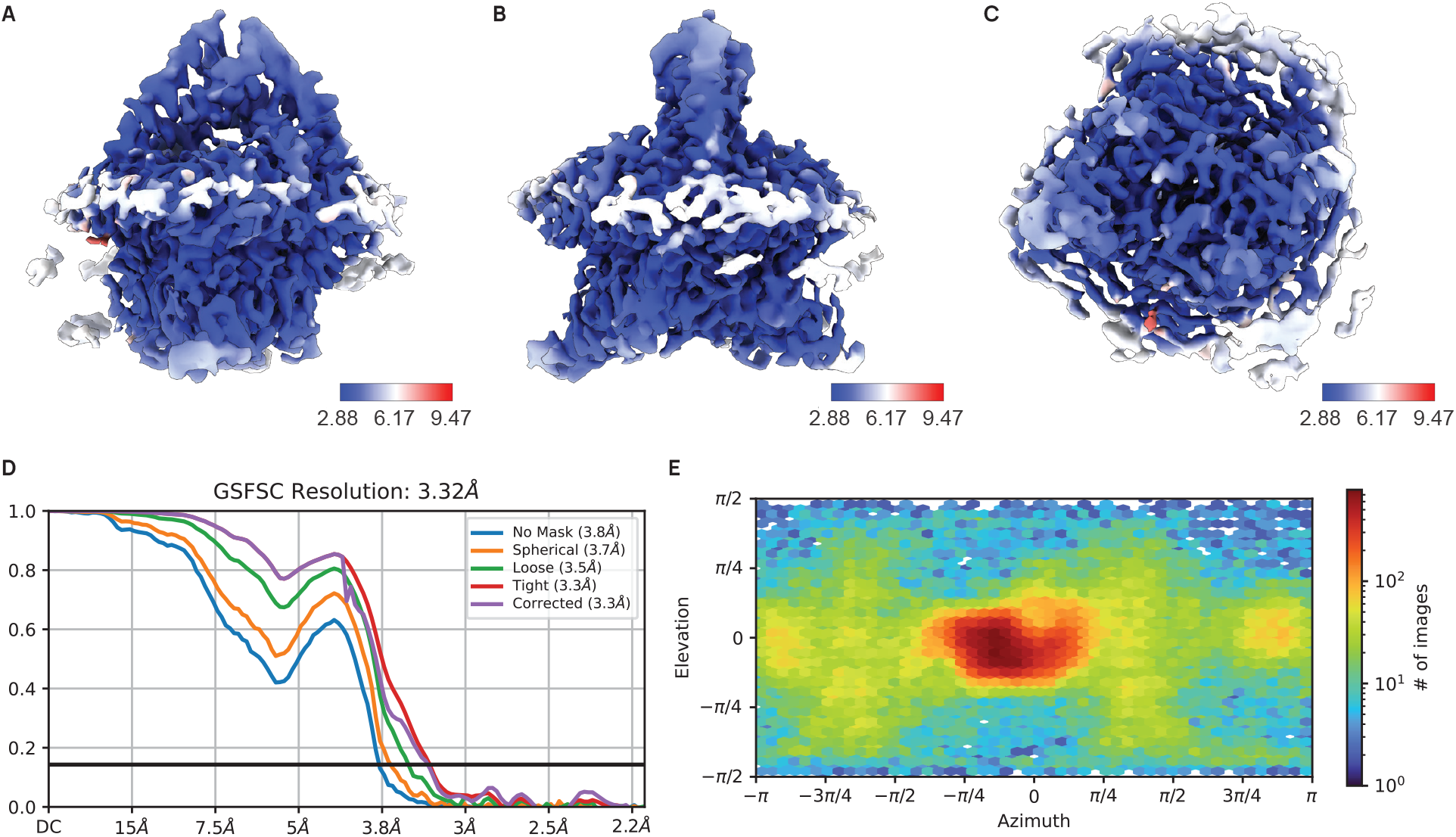
Cryo-EM structure validation for TASK-1 at pH 6.0. Final sharpened map colored by local resolution and viewed from (A) the membrane plane (B) rotated 90° (C) the cytoplasmic side. (D) Fourier shell correlation (FSC) versus resolution between half maps from final refinement. (E) View angle distribution of final particle stack.

**Figure S2.**
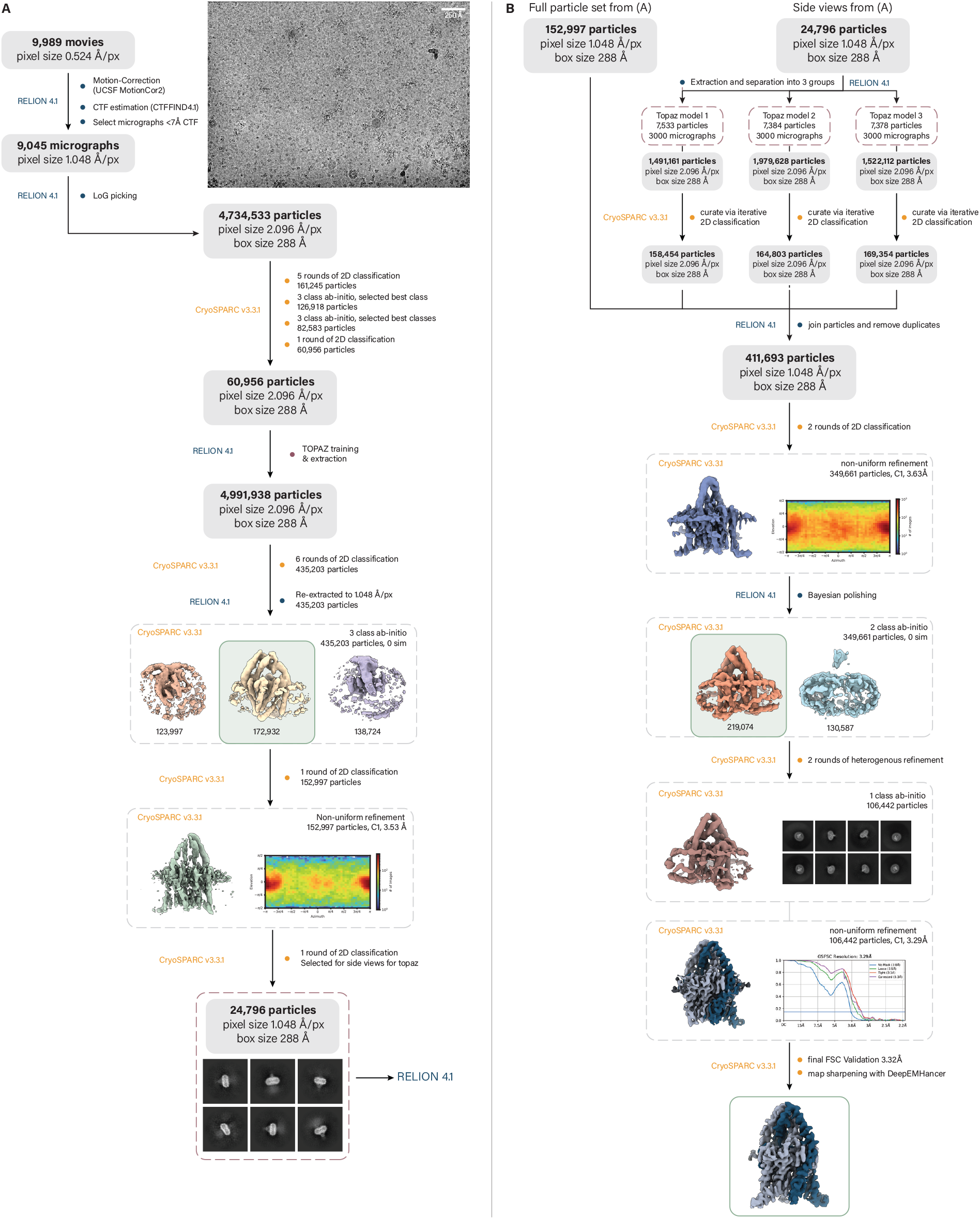
Cryo-EM data processing pipeline for TASK-1 at pH 6.0. (A) Representative micrograph and initial cryo-EM data processing. (B) Cryo-EM data processing pipeline approach for finding additional side views of the channel to alleviate observed anisotropy with representative 2D class averages, reconstructed map from final particle stack, and DEEPEMHANCER sharpened map.

**Figure S3.**
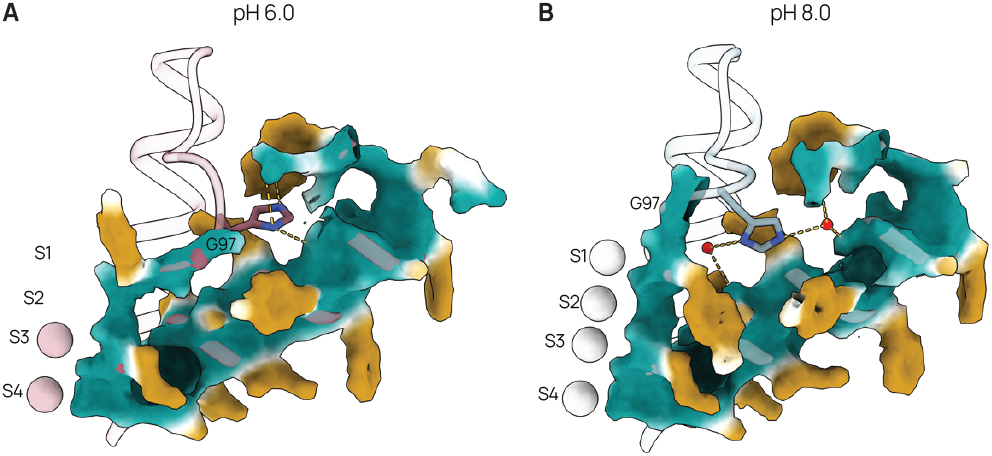
Hydrophobicity surface map of H98 pocket at pH 6.0 and pH 8.5. Representative hydrophobicity structure surface map of the H98 pocket at (a) pH 6.0 and (b) pH 8.5. At pH 6, H98 is pulled away from the channel conduction axis toward W78, resulting in an upward shift of Y96 and the movement of G97 into the cavity. At pH 8.0, G97 and H98 are shifted back toward the conduction axis of the channel and Y96 is rotated around toward the cavity, generating a hydrophobic pocket for H98.

**Figure S4.**
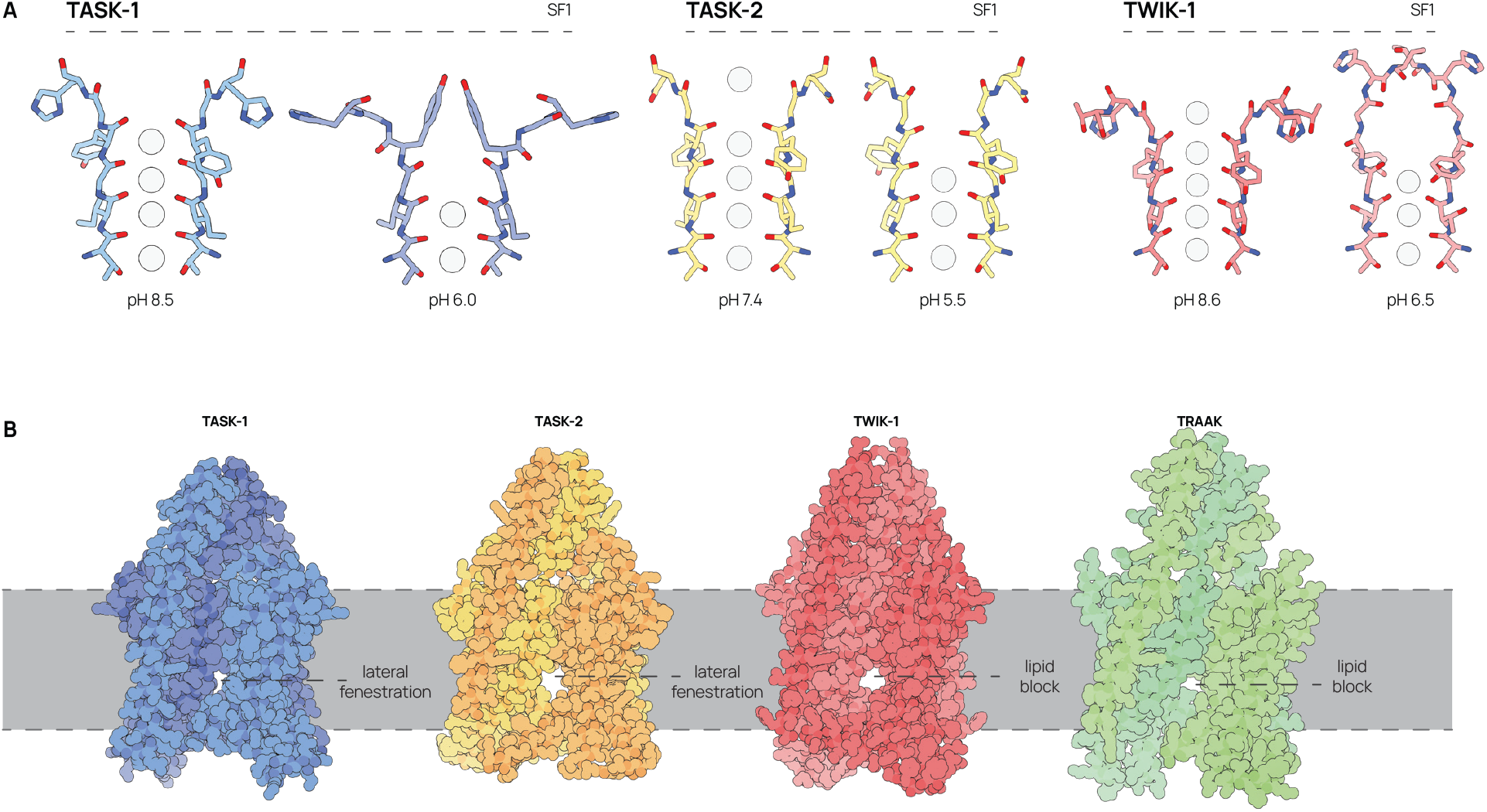
Comparison of K2P selectivity filter conformational changes in response to pH. (A) Structures of the selectivity filters of pH-sensitive K2Ps TASK-1, TASK-2 and TWIK-1 at high and low pH. (B) cryo-EM models of TASK-1, TASK-2 (PDB: 6WLV), TWIK-1 (PDB: 7SK1), and TRAAK (PDB: 4WFF) from the membrane plane. All four channels display a lateral fenestration between TM2 and TM4 below the selectivity filter. To date, TWIK-1, TWIK-2 and TRAAK channels have been proposed to have gating states stabilized by the insertion of inner leaflet lipid acyl chains into the channel cavity through this fenestration.

